# A novel mosaic tetracycline resistance gene *tet*(S/M) detected in a multidrug-resistant pneumococcal CC230 lineage that underwent capsular switching in South Africa

**DOI:** 10.1101/718460

**Authors:** Stephanie W. Lo, Rebecca A. Gladstone, Andries J. van Tonder, Mignon du Plessis, Jennifer E. Cornick, Paulina A. Hawkins, Shabir A. Madhi, Susan A. Nzenze, Rama Kandasamy, KL Ravikumar, Naima Elmdaghri, Brenda Kwambana-Adams, Samanta Cristine Grassi Almeida, Anna Skoczynska, Ekaterina Egorova, Leonid Titov, Samir K. Saha, Metka Paragi, Dean B. Everett, Martin Antonio, Keith P. Klugman, Yuan Li, Benjamin J Metcalf, Bernard Beall, Lesley McGee, Robert F. Breiman, Stephen D. Bentley, Anne von Gottberg, on behalf of The Global Pneumococcal Sequencing Consortium

## Abstract

**Objective:** We reported a novel tetracycline-resistant gene in *Streptococcus pneumoniae* and investigated its temporal spread in relation to nationwide clinical interventions.

**Methods:** We whole genome sequenced 12,254 pneumococcal isolates from twenty-nine countries on an Illumina HiSeq Sequencer. Serotypes, sequence types and antibiotic resistance were inferred from genomes. Phylogeny was built based on single-nucleotide variants. Temporal changes of spread were reconstructed using a birth-death model.

**Results:** We identified *tet*(S/M) in 131 pneumococcal isolates, 97 (74%) caused invasive pneumococcal diseases among young children (59% HIV-positive, where HIV status was available) in South Africa. A majority of *tet*(S/M)-positive isolates (129/131) belong to clonal complex (CC)230. A global phylogeny of CC230 (n=389) revealed that *tet*(S/M)-positive isolates formed a sub-lineage that exhibited multidrug-resistance. Using the genomic data and a birth-death model, we detected an unrecognised outbreak of this sub-lineage in South Africa between 2000 and 2004 with an expected secondary infections (R) of ~2.5. R declined to ~1.0 in 2005 and <1.0 in 2012. The declining epidemic coincided and could be related to the nationwide implementation of anti-retroviral treatment (ART) for HIV-infected individuals in 2004 and PCVs in late 2000s. Capsular switching from vaccine serotype 14 to non-vaccine serotype 23A was observed within the sub-lineage.

**Conclusions:** The prevalence of *tet*(S/M) in pneumococci was low and its dissemination was due to an unrecognised outbreak of CC230 in South Africa prior to ART and PCVs. However, capsular switching in this multidrug-resistant sub-lineage highlighted its potential to continue to cause disease in the post-PCV13 era.

## Introduction

*Streptococcus pneumoniae* is a major bacterial cause of disease in young children. Since 2000, pneumococcal conjugate vaccines (PCVs) targeting up to 13 serotypes were gradually introduced into childhood immunisation programmes in many countries and have significantly reduced pneumococcal deaths globally by 51% and 75% in HIV-uninfected and HIV-infected children aged <5 years, respectively, resulting in saving an estimated 375,000 lives annually when compared with the estimated mortality rate in the pre-vaccine era ^1^. However, increasing invasive pneumococcal disease (IPD) caused by non-vaccine serotype pneumococci has been observed in numerous locations including England and Wales ^2^, France ^3^, Germany ^4^ and Israel ^5^, a phenomenon known as serotype replacement. Serotype replacement could be mediated by capsular switching, in which a *cps* locus encoding vaccine-type (VT) capsule is replaced by a *cps* locus encoding non-vaccine-type (NVT) capsule through homologous recombination ^6^. Capsular switching within multidrug-resistant lineages, especially those recognised by the Pneumococcal Molecular Epidemiology Network (PMEN, http://spneumoniae.mlst.net/pmen/pmen.asp), is of increasing concern, as these expansions can reduce overall vaccine effectiveness in preventing IPD and temper the reduction in antimicrobial-resistant pneumococcal infections associated with introduction of PCVs ^7^. The persistence of the multidrug resistant lineage ST156 (Spain ^9V^-3, PMEN3) in the USA following the introduction of PCV13 provides a clear example of a historically successful lineage that underwent a capsular switch from VT (serotype 9V, 14 and 19A) to NVT (serotype 35B) and continued to cause IPD in the post-vaccine era ^7–9^.

Resistance to tetracycline has been frequently observed in *S. pneumoniae* ^10^. The genetic basis was shown to be the *tet*(M), less commonly *tet*(O), which encode for a ribosomal protection protein that prevents tetracycline binding to the bacterial 30S ribosome subunit ^10, 11^. Eleven other classes of ribosomal protection proteins such as *tet*(S) and twelve mosaic structure of *tet* genes such as *tet*(S/M) have not been found previously in pneumococci (http://faculty.washington.edu/marilynr/). The *tet*(S), originally discovered in *Listeria monocytogenes* strain BM4210 ^12^, was occasionally found in a variety of streptococci, including *S. suis* (NCBI accession number KX077886) ^13^, *S. infantis* (NCBI accession number JX275965), and *S. dysgalactiae* (NCBI accession number EF682210) ^14^ and is associated with a transposase-containing element *IS1216*, which potentially mediates chromosomal rearrangement. The mosaic *tet*(S/M) was observed on a Tn*916* element in *S. intermedius* ^15^ and an *IS1216* composite in *S. bovis* ^16^. Using a dataset of 12,254 pneumococcal genomes from The Global Pneumococcal Sequencing (GPS) project (https://www.pneumogen.net/gps/), we identified a novel genetic basis for tetracycline resistance *tet*(S/M) in *S. pneumoniae* and characterised its genetic background in relation to nationwide clinical interventions.

## Material and Methods

### Isolate collection

In the GPS project, each participating country randomly selected disease isolates collected via laboratory-based surveillance and carriage isolates via cohort-studies using the following criteria: ~50% isolates were from children ≤ 2 years, 25% from children 3-5 years, and 25% from individuals >5 years. By May 2017 (last accessed to the GPS database for this study), 12,254 isolates, representing 29 countries, in Africa (65%), North America (14%), Asia (9%), South America (8%), and Europe (4%), were sequenced, passed quality control and included in this study. The collection spanned 26 years between 1991 and 2016 and included both carriage (n=4,863) and disease isolates (n=7,391). We compiled the metadata including age, year of collection, sample source, HIV status and phenotypic antimicrobial susceptibility testing results, where available, from each participating site. In children < 18 months of age, HIV status was confirmed by PCR assay. MIC results were interpreted according to Clinical Laboratory Standards Institute M100-S24 ^17^ When MIC was analysed as “>X”, MIC was approximated as value 2X for median and interquartile range calculations.

### Genome sequencing and analyses

The pneumococcal isolates were whole genome sequenced on an Illumina HiSeq platform and raw data were deposited in the European Nucleotide Archive (ENA) (Supplementary metadata). We inferred serotype, multilocus sequence types (MLSTs) and resistance profile for penicillin, chloramphenicol, cotrimoxazole, erythromycin and tetracycline from the genomic data as previously described ^18^. The *tet*(S/M) gene was identified with a *tet*(S/M) reference sequence (NCBI accession number AY534326) using ARIBA ^19^.

To reconstruct a global phylogeny, an additional collection of CC230 isolates (n=130) from previous studies ^20–24^, together with the CC230 collection (n=259) in the GPS dataset were included. The phylogeny was built as previously described ^18^. Based on the international genomic definition of pneumococcal lineages, all CC230 isolates in this study belong to Global Pneumococcal Sequence Cluster (GPSC)10 ^18^. The metadata and analysis results of CC230 can be interactively visualized online using the Microreact tool at https://microreact.org/project/GPS_tetSM.

### Temporal changes of *tet*(S/M) CC230 sub-lineage

Coalescent analysis was performed on *tet*(S/M) CC230 sub-lineage (n=129) to date the most recent common ancestor (MRCA) and reconstruct the population demographic history. First, we tested the presence of temporal signal by a linear regression of root-to-tip distances against year of collection using TempEst v1.5 ^25^. Next, a timed phylogeny was constructed using BEAST v2.4.1 ^26^. The Markov chain Monte Carlo (MCMC) chain was run for 100 million generations, sampled every 1000 states using the general time-reversible (GTR) model of nucleotide substitution and the discrete gamma model of heterogeneity among sites. Finally, the population demographic history was reconstructed using a birth-death model ^27^ to examine the temporal changes with the *tet*(S/M) CC230 sub-lineage invasive isolates (n=105) but not carriage isolates (n=24), because the model assumes that once an individual is diagnosed with IPD, the individual is no longer transmitting due to treatment and recovery, death, or being socially removed from susceptible individuals. Thus, it appeared to be logical to apply this model to the disease but not carriage isolates. This model overcomes the limitations of the coalescent-based skyline plot and is able to examine whether introduction of an intervention had an impact on the epidemiological dynamics in a bacterial population ^28^. The birth-death skyline plot shows the effective reproductive number (R) over time. R is defined as the number of expected secondary infections of an infected individual. R>1 indicates a growing epidemic, whereas R<1 indicates a declining epidemic. Notably, R≥1 can be reflected in the coalescent-based skyline plot analysis, whereas R<1 cannot. Therefore, we expected the birth-death skyline model would be a better fit for our data. Other Bayesian population size models (coalescent constant, coalescent exponential and Bayesian skyline) in combination with strict and lognormal-relaxed molecular clocks were also applied for comparisons using BEAST.

### Integrative and conjugative element (ICE)

The ICE was extracted from the *de novo* assemblies of CC230 isolates and compared using EasyFig version 2.2.2. The NCBI accession numbers for the representative ICE sequences in Figure 5 were FM211187 (ICESp23FST81), MH283017 [ICES*p*14ST230 with *tet*(M)], MH283012 [ICES*p*14ST230 with *tet*(S/M) and omega cassettes], MH283013 [ICES*p*14ST230 with *tet*(M) and Omega], MH283012 [ICES*p*14ST230 with *tet*(M) and Tn*917*], MH283016 [ICES*p*19AST2013], MH283015 [ICES*p*17FST8812], and MH283014 [ICES*p*14ST156].

## Results

### Prevalence of tet(S/M) in a global collection of S. pneumoniae

A novel tetracycline-resistant gene *tet*(S/M) was identified in 131 pneumococcal isolates (1%, 131/12,254) from South Africa (n=123), Malawi (n=5), and one each from Brazil, Mozambique, and the USA. They were isolated from sterile body sites (invasive isolates): blood (n=73), cerebrospinal fluid (n=30), pleural fluid (n=4), and from the nasopharynx (carriage isolates) (n=24). In South Africa, *tet*(S/M) was found in 3.5% (103/2920) of the invasive isolates that were submitted to the GPS project from 2005-2014 and 1.2% (20/1701) carriage isolates that were collected in Agincourt and Soweto between 2009 and 2013. Of the 103 invasive isolates, 94% (97/103) were from children with IPD aged ≤ 5 years (Fig. S1). HIV status was known in only 44% (54/123) of individuals with *tet*(S/M)-positive pneumococci; 59% (32/54) were HIV-positive, in which 94% (30/32) were children aged ≤ 5 years.

Among the *tet*(S/M)-positive isolates, the minimum inhibitory concentration (MIC) to tetracycline was determined by either E-test (n=56) or broth dilution (n=48). E-test showed a median MIC of 8 mg/L with an interquartile range of 6-9 mg/L and 16 mg/L by broth dilution. Based on the CLSI guideline, 99% (103/104) and 1% (1/104) were fully and intermediately resistant to tetracycline, respectively. The *tet*(S/M) in this study showed 100% nucleotide identity, except for one isolate (GPS_ZA_1982) from South Africa which varied from the others at G1769A and resulted in a substitution R590Q. This isolate remained resistant to tetracycline with a MIC of >8 mg/L when measured by broth dilution. Unlike the two previously reported *tet*(S/M) alleles from *S. intermedius* ^15^ and *S. bovis* ^16^, the amino acid sequence of *tet*(S/M) in this study showed 100% identity to Tet(S) (NCBI accession number FN555436) across the first 613 amino acids, with the final 32 amino acids at the C-terminus end being identical to Tet(M) (NCBI accession number EFU09422) (Figure 1). Examining the promoter regions revealed that the −10 (TATTAT) and −35 (TTTACA) promoter sequence was of *tet*(M) origin, rather than *tet*(S) origin. Between the promoter region and start codon of *tet*(S/M), a 38-bp stem loop which is potentially involved in transcriptional regulation ^29^ was found in all *tet*(S/M) genes (Figure 1), apart from one disease isolate (GPS_ZA_1926) from South Africa. The deletion did not affect the tetracycline resistance level, as the MIC remained at >8 mg/L when measured by the broth dilution method.

**Figure 1.**
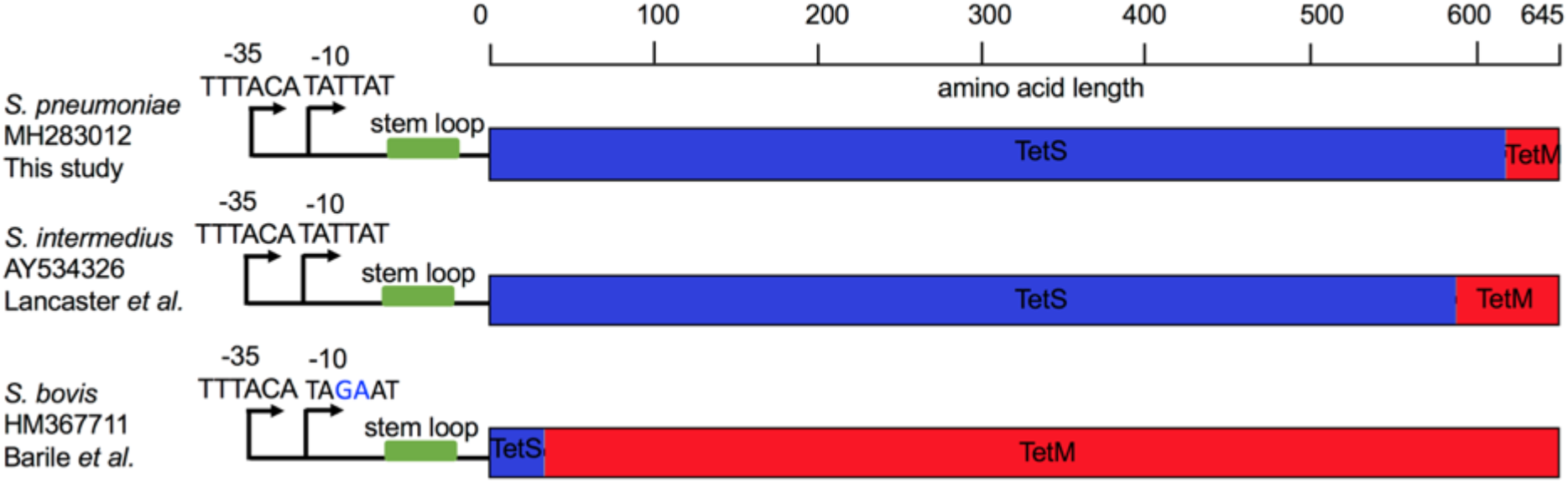
Schematic representation of the mosaic structure of *tet*(S/M) alleles of the current and previous studies. The reference sequences for *tet*(M) and *tet(S)* were retrieved from NCBI Genbank using accession number EFU09422 and FN555436, respectively.

### Phylogeny and characteristics of *tet*(S/M) CC230 sub-lineage

All 131 *tet*(S/M)-positive isolates belonged to CC230, except for one Brazilian and one Malawian isolate belonging to CC156 and ST5359 (a singleton not belonging to any CC), respectively. The global CC230 phylogeny showed that all *tet*(S/M)-positive isolates formed a sub-lineage which predicted to be resistance to penicillin, erythromycin, tetracycline and cotrimoxazole (Figure 2). The *tet*(S/M) sub-lineage was associated predominantly with VT 14 (98%,127/129) but was also found in two NVT 23A isolates. The two serotype 23A isolates, which both belonged to ST11106 (a single-locus variant of ST230), were recovered from infants after the introduction of PCV13. One was isolated from a nasopharyngeal sample in Soweto in 2012 and the other from blood culture in Johannesburg in 2014. The serotype 23A *cps* locus sequences of these two isolates were identical, and their *cps*-flanking *pbp* loci (*pbp*1A and *pbp*2X) were also identical to the majority of the serotype 14 isolates within the *tet*(S/M) sub-lineage, exhibiting resistance to penicillin with MIC of 2 mg/L. To identify the potential donor of the serotype 23A *cps* locus, a phylogenetic tree was built using the *cps* sequences from all serotype 23A (n=130, belong to eight lineages) pneumococci in the GPS database. This analysis showed that the serotype 23A *cps* loci of these two CC230 isolates clustered with those originating from a serogroup 23 lineage GPSC7, which is predominantly (99%, 145/146) represented by CC439 (Figure S2), with pairwise nucleotide similarity of 99.97% (24,818/24,825) and 100% coverage. The seven nucleotide variations were found within the *IS630* transposase downstream of *dexB*.

**Figure 2.**
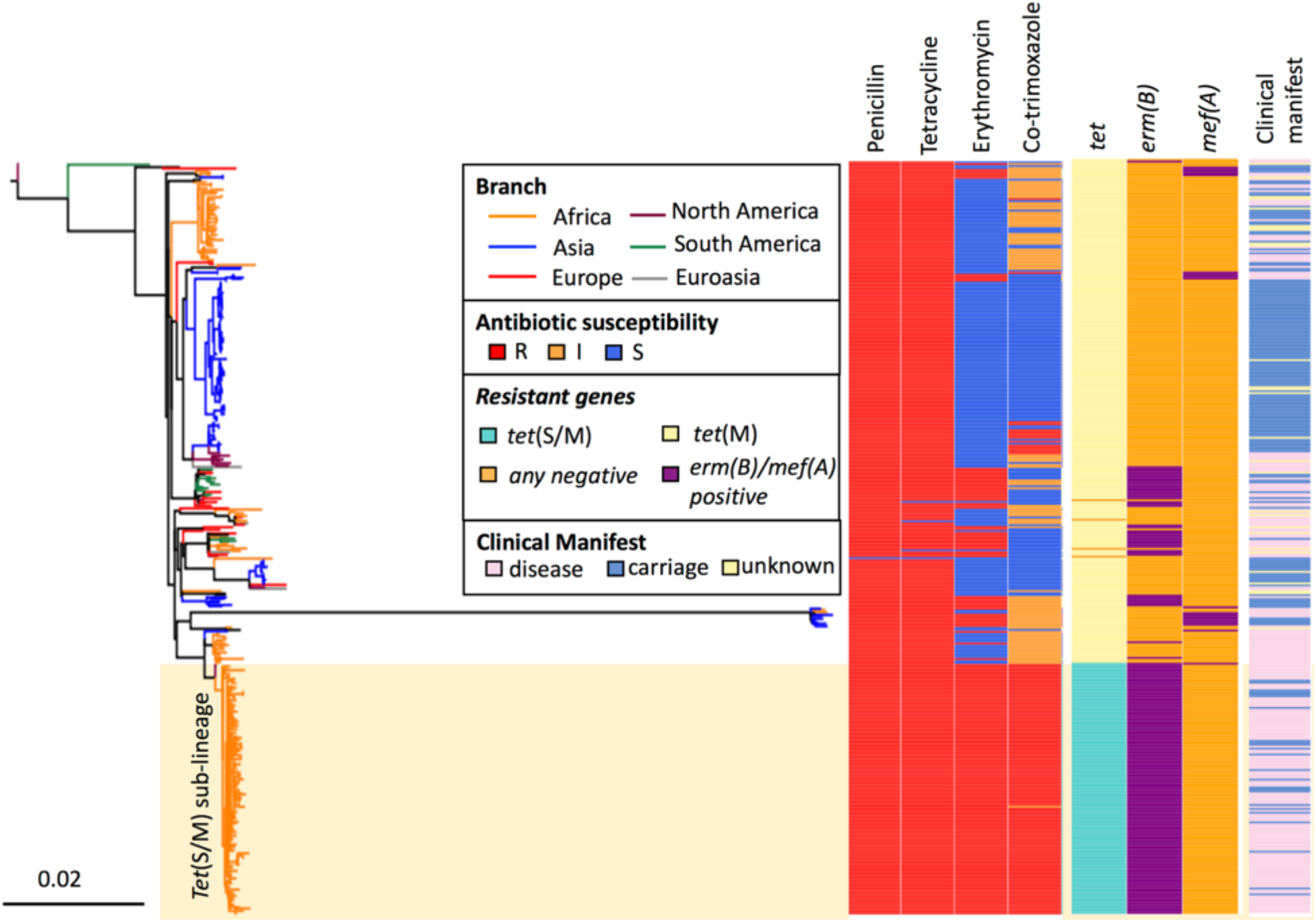
A SNP tree constructed with CC230 *tet*(S/M)-positive isolates (n=129) and *tet*(S/M)-negative carriage/disease isolates (n=260) collected from twenty countries. The tree was built based on 13,405 SNPs extracted from an alignment outside recombination regions, created by mapping reads of each isolate to the sequence of a ST230 reference strain, PMEN global clone Denmark^14^-32, PMEN32 (ENA accession number ERS1706837). Penicillin resistance was predicted based on the *pbp*1a, *pbp*2x, *pbp*2b sequences (1, 2); tetracycline and erythromycin resistance were predicted based on the presence of *tet*(M) and *tet*(S/M), and *erm*(B) and *mef*(A), respectively. Cotrimoxazole resistance was predicted based on the presence of mutation I100L in *folA* and any indel within amino acid residue 56-67 in *folP* while presence of either mutation predicted to confer cotrimoxazole-intermediate phenotype.

### Temporal spread of the tet(S/M) CC230 sub-lineage

The sub-lineage showed a temporal signal in terms of SNP accumulation against time (R^2^ = 0.4094, p value = 0.001; Figure S3). Using a birth-death model in BEAST, the *tet*(S/M) sub-lineage was estimated to emerge around 1994 (95% highest posterior density [HPD]: 1991-1996); the MRCA for the African clade was 1998 (95% HPD: 1996-2000) and for the two serotype 23A isolates was 2009 (95% HPD: 2007-2011) (Figure 3). The temporal changes of spread were reconstructed based on a birth-death skyline plot and coalescent-based skyline plot (Figure 4). Both skyline plots showed that the *tet*(S/M) sub-lineage expanded at the beginning of the year 2000 and growth continued until around 2004. The decline of the *tet*(S/M) sub-lineage was only captured by the birth-death skyline plot in or around 2005, from expected secondary infections (R) of ~2.5 to ~1, and steadily declined until 2012 when the median and HPD of R were below one, indicating a declining epidemic. The coalescent-based skyline plot failed to detect the impact of the epidemic decline as described in a previous study ^27^.

**Figure 3.**
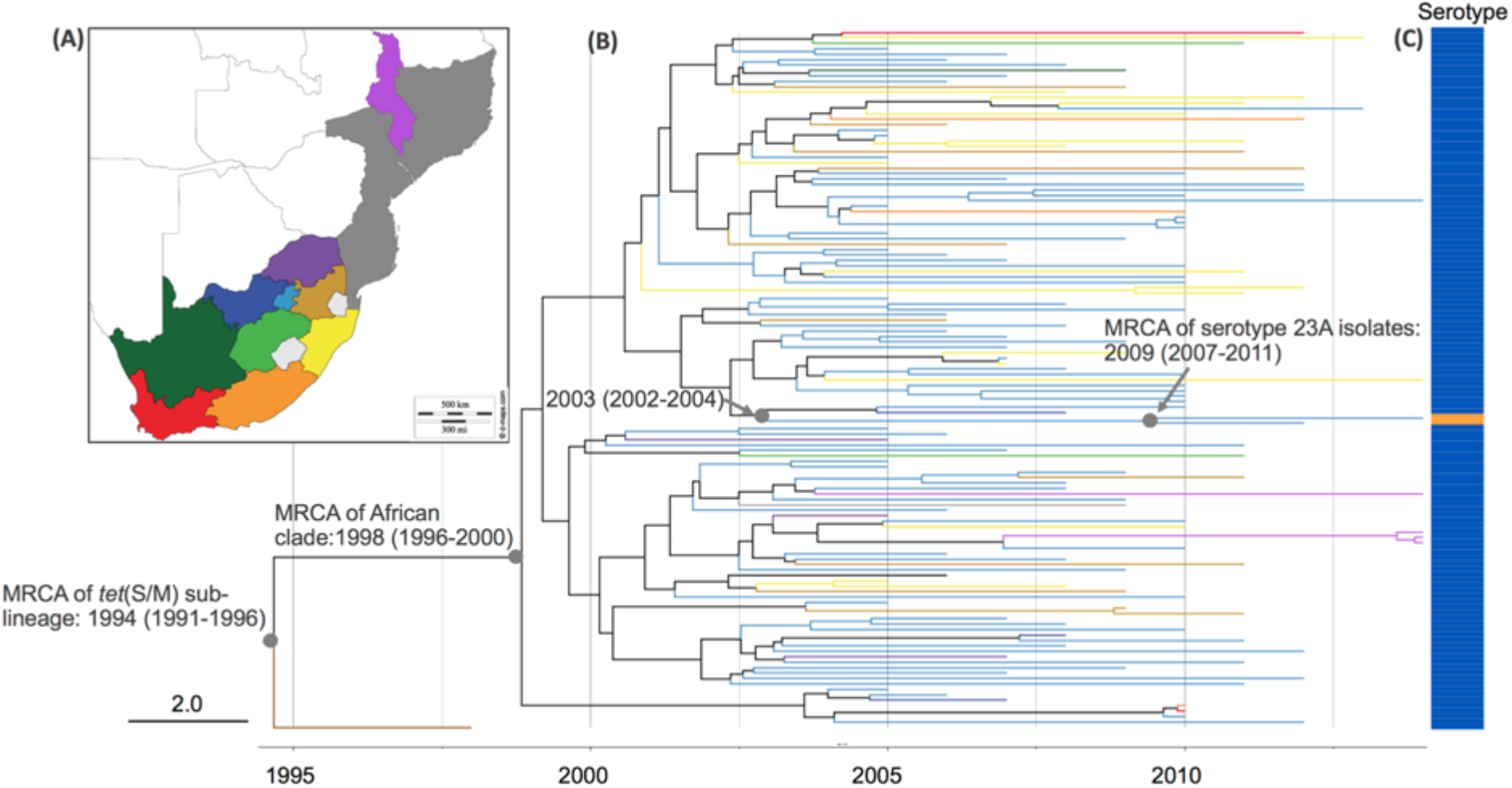
(A) Malawi, Mozambique and administration regions of South Africa (B) Timed phylogeny for *S. pneumoniae tet*(S/M) CC230 sub-lineage (n=129) reconstructed using BEAST. Tree branches are coloured according to the geographical locations in (A), except for the branch for an isolate collected from the United States coloured in brown. (C) Vaccine serotype 14 is indicated in blue, whereas non-vaccine serotype 23A in orange.

**Figure 4.**
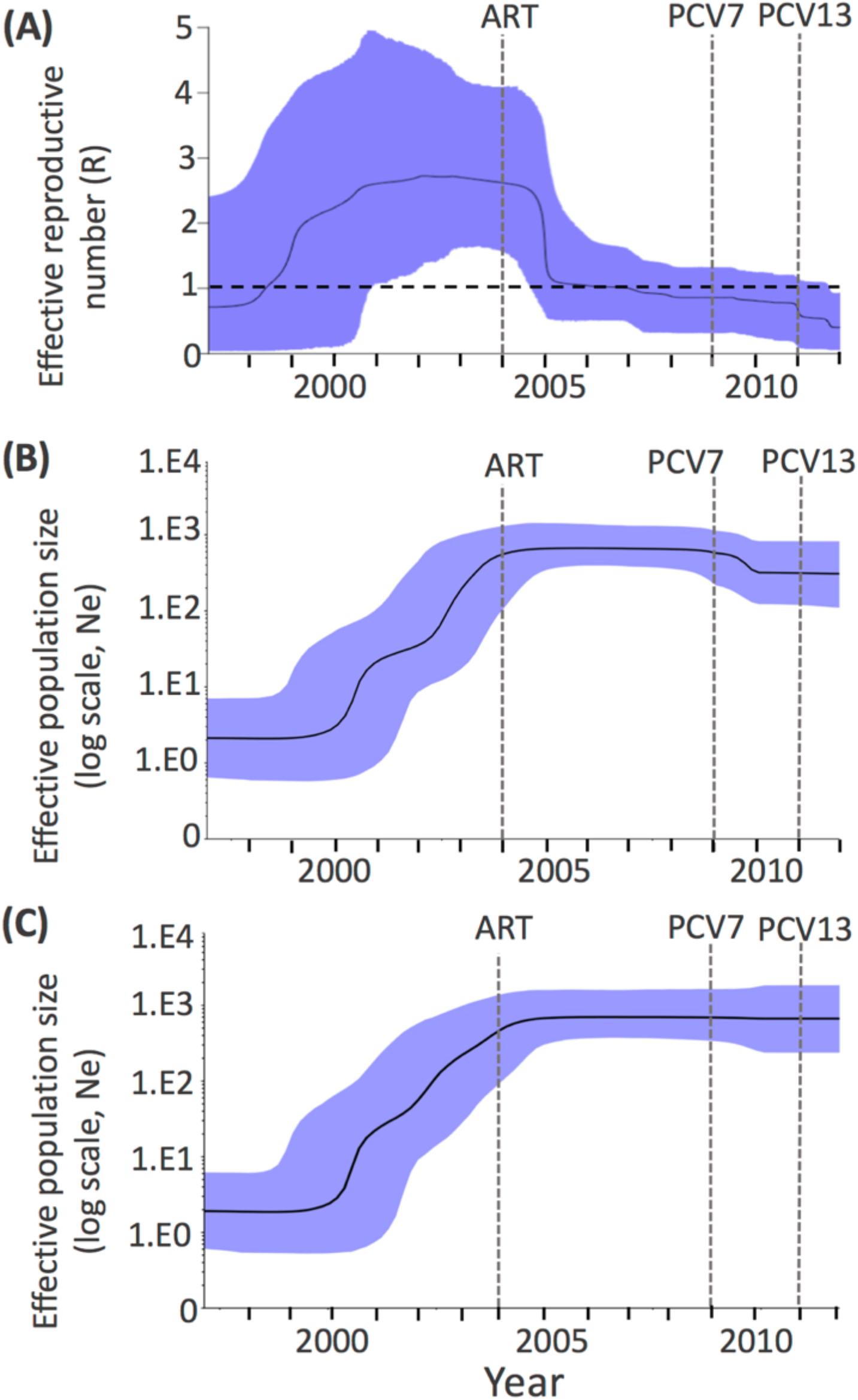
(A) Birth-death skyline plot of inferred changes in effective reproductive number (R) of *S. pneumoniae tet*(S/M) CC230 sub-lineage using IPD isolates (n=105). (B) Coalescent-based skyline plot of inferred changes in the effective population size (Ne) of *S. pneumoniae tet*(S/M) CC230 sub-lineage using both IPD and carriage isolates (n=129) and (C) using only IPD isolates (n=105). The black solid line shows the median of R in (A) and Ne in (B) and (C), respectively. The background area represents the 95% highest posterior density intervals. R>1 indicates a growing epidemic, whereas R<1 indicates a declining epidemic. ART, antiretroviral treatment; PCV7, seven-valent pneumococcal conjugate vaccine; PCV13, thirteen-valent pneumococcal conjugate vaccine.

### ICE carrying tet(S/M)

The acquisition of tetracycline and erythromycin resistance determinants by CC230 was the result of the insertion of a Tn5253-type ICE, which shared a similar structure to ICESp23FST81 identified in PMEN1 (Figure 5). Both the *tet*(M) or *tet*(S/M) genes detected in this study were carried on a conserved conjugative Tn*916* transposon (Figure S4). Of the 172 macrolide-resistant isolates, insertions of either the ‘omega’ element (n=165) or Tn917 (n=7) harbouring *erm*(B) were found up- or downstream of the *tet* gene, respectively (Figure 5). The insertion of the ‘omega’ element truncated the gene encoding the replication initiation factor, creating an 8-bp direct repeat, CAAAAAAA. The insertion of Tn917 disrupted the gene *orf9* which encodes a putative conjugative transposon regulator. No direct repeats were found.

**Figure 5.**
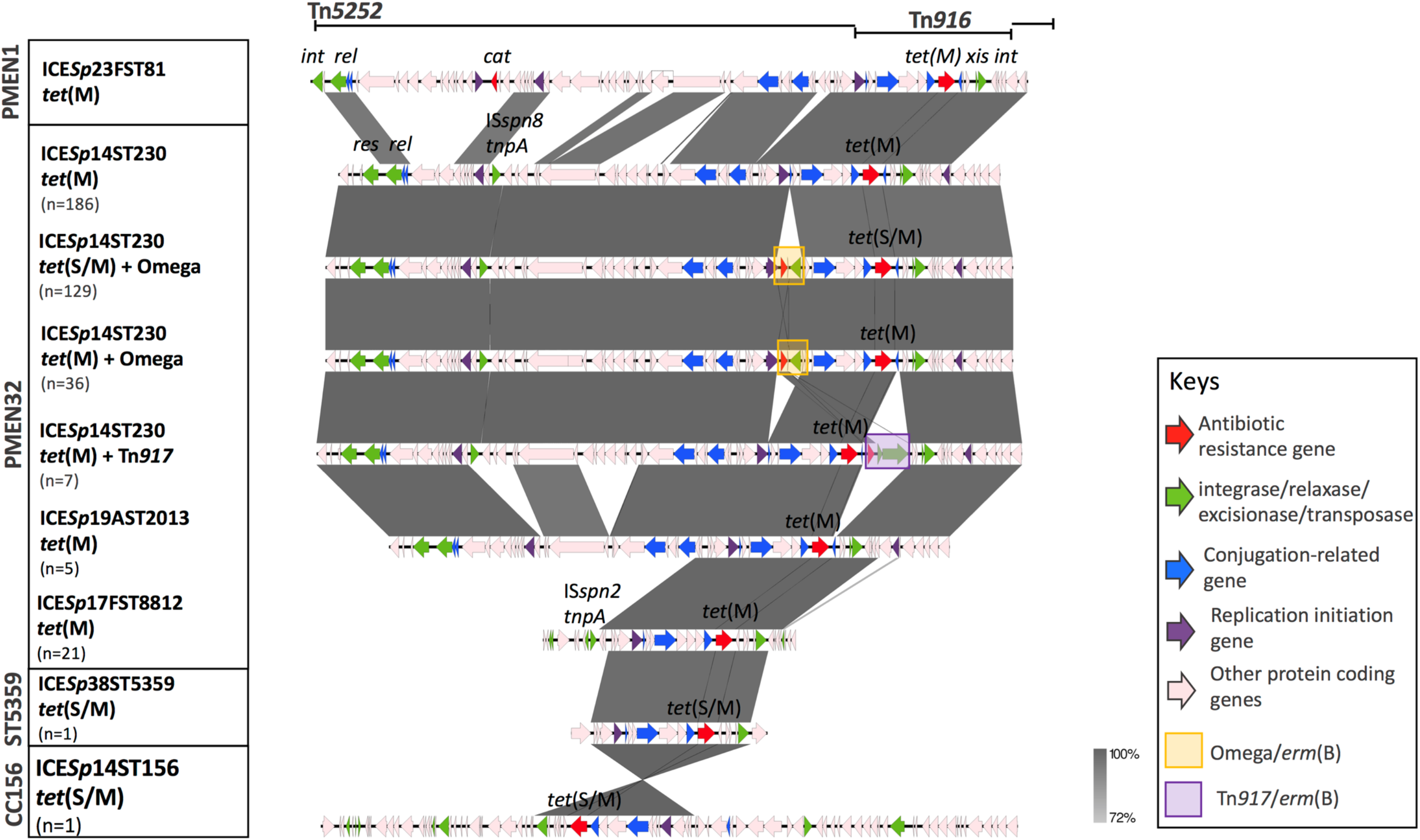
Comparison of integrative and conjugative element (ICE) identified in clonal complex (CC)230 with Spain^23F^-1 (PMEN1). Grey bands between the sequences indicate BLASTN matches.

## Discussion

We used WGS to identify a novel mosaic structure of *tet*(S/M) in *S. pneumoniae*. This approach overcame the limitation of PCR that requires specific primers to detect known antibiotic resistance genes. Compared with *tet*(M), the prevalence of *tet*(S/M) was low. They were mainly found in a CC230 sub-lineage that predominantly expressed VT 14 and exhibited multidrug resistance in South Africa. Together with its conserved nucleotide sequence and genomic location, our finding strongly suggested a clonal expansion of *tet*(S/M)-positive CC230 isolates within South Africa prior to the introduction of PCVs.

The convergence of antimicrobial resistance and virulence in the CC230 sub-lineage probably contributed to its expansion in disease-causing populations prior to the introduction of PCVs. Unlike what was observed in other countries, CC230 in South Africa predominantly expressed a highly invasive serotype 14 capsule ^18^ and was the clone that represented most of the serotype 14 isolates (43%) in the pre-vaccine era, when serotype 14 was the most prevalent serotype causing IPD in South Africa ^30^. Any controlling measure to decrease this lineage would not only result in a reduction in the IPD burden but also multidrug-resistant IPD incidence.

The birth-death model estimated that the decline of the *tet*(S/M) CC230 sub-lineage started around 2005, one year after the national ART programme was launched to treat HIV-infected individuals in South Africa ^31^. The prediction was consistent with a 41% reduction of IPD incidence among HIV-infected children after the introduction of ART ^31^. Among the IPD caused by *tet*(S/M) CC230 isolates, almost 60% occurred in HIV-positive children. This observational evidence strengthened that ART was likely to contribute to the decline before PCV introduction. In contrast, the large-scale use of cotrimoxazole as prophylaxis to prevent bacterial infections among HIV-positive individuals was unlikely to be responsible for the decline, as the sub-lineage was resistant to cotrimoxazole. The further decline in 2012 predicted by the model echoed the epidemiological finding that IPD caused by VT pneumococci significantly decreased among children in 2012 ^32^. Our finding demonstrated that we could effectively reconstruct the temporal spread of an epidemic using genomic data and highlighted the possible use of routine genomic surveillance to identify outbreaks as they occur in the future.

The MRCA of two CC230 isolates expressing NVT 23A was dated to emerge around 2009, the year when PCV7 was introduced. However, the long branch leading to the MRCA from the internal node that shared with the closely related serotype 14 isolates indicated that the window of time for capsular switching could be between 2002 and 2011. Although the invasive disease potential for serotype 23A is low ^18^, a significant increase of this serotype in IPD cases was reported from England ^2, 33^, Stockholm ^34^ and Taiwan ^35^ after the implementation of PCV13. Serotype 23A is primarily associated with CC338 (GPSC5, PMEN26) and CC439 (GPSC7), and is thus rarely found in a CC230 genetic background. Such serotype and genotype combination were only identified in two ST9396 isolates (single locus variant of ST230) from China in 2013 and one ST10921 isolate (double locus variant of ST230) from Poland in 2013 in the MLST database. In South Africa, CC439 accounted for 62% of serotype 23A (both carriage and disease) isolates ^18^ is the potential donor of the serotype 23A *cps* to the *tet*(S/M) CC230 sub-lineage, highlighting that capsular switching with the prevalent NVT lineage could enable a VT lineage to evade the vaccine. Capsular switching is usually a result of homologous recombination. When compared with other 620 GPSCs, GPSC10 which included 98% (258/262) of CC230 isolates is a very recombinogenic lineage which had a significantly high recombination rate [GPSC10 r/m: 10.9 vs median of 35 dominant GPSCs: 8.3 (1^st^ – 3^rd^ quartile, 5.7-10.7) p value < 0.0001, Wilcoxon signed-rank test] ^18^. Given this recombinogenic nature, together with the established multidrug resistant genotypes, it is of concern that any further capsular switching may increase the chance of this multidrug-resistant lineage surviving and continuing to cause invasive disease.

Like *tet*(M), *tet*(S/M) was also carried by a highly mobile conjugative transposon, Tn*916*, with a broad host range. Conserved genetic environment of *tet*(M) and *tet*(S/M) indicates that the recombination resulting in the mosaic structure of *tet*(S/M) probably occurred after the acquisition of the gene by Tn*916*. Comparison of *tet*(M) sequences in the current collection also revealed a high degree of allelic variations that were probably due to homologous recombination ^36^. This finding is consistent with previous studies which suggest that the *tet* evolved separately from Tn*916* ^10, 36^. However, the driving force behind the evolution of *tet* genes remains unclear, given that tetracycline is not used as a first-line antibiotic to treat pneumococcal disease and was seldom used in young children ^37^. The allelic diversity of *tet* gene may be maintained by 1) frequent recombination among *S. pneumoniae* and with closely related species such as normal nasopharyngeal resident *S. mitis* ^38^ and zoonotic pathogen *S. suis* ^39^; 2) antibiotic-selective pressure via food chain, as tetracycline is widely used in agriculture ^40^ and its residue is detected in milk ^41^. Future studies that investigate the driving force behind will improve our understanding to develop preventive measure to reduce tetracycline resistance in *S. pneumoniae*.

In conclusion, we identified a novel tetracycline-resistant determinant *tet*(S/M) in *S. pneumoniae* and showed that its dissemination is due to a clonal expansion of the multidrug-resistant lineage CC230 in South Africa where the HIV burden is high. With genomic data, we successfully detected the declines in transmission of this multidrug-resistant lineage using a birth-death model, and the fall of this lineage may correlate to the improved treatment of HIV-infected individuals and the implementation of PCVs. Capsular switching within this lineage is potentially of public health importance and may erode the beneficial effect brought about by the implementation of PCVs. The capacity for continuous genomic surveillance in the post-vaccine era provides critical opportunities for monitoring and forecasting the rise of multidrug-resistant pneumococcal lineages that may also undergo vaccine evasion through capsular switching events.

## Acknowledgements

We deeply appreciate all members of the GPS consortium for their collaborative spirit and determination for the monumental task of sampling, extracting data, and for their intellectual input to this manuscript. We extend our special thanks to Linda De Gouveia from National Institute for Communicable Diseases of the National Health Laboratory Service, South Africa who helped us with the antimicrobial susceptibility testing. We appreciated critiques and suggestions from all team members in the Genomics of Pneumonia and Meningitis (and neonatal sepsis) group and technical support from the Pathogen Informatic Team in the Parasites and Microbe Programme at the Wellcome Sanger Institute.

Members of The Global Pneumococcal Sequencing Consortium:

Abdullah W Brooks, Alejandra Corso, Alexander Davydov, Andrew J Pollard, Anna Skoczynska, Betuel Sigauque, Deborah Lehmann, Diego Faccone, Elena Voropaeva, Eric Sampane-Donkor, Ewa Sadowy, Godfrey Bigogo, Helio Mucavele, Houria Belabbès, Idrissa Diawara, Jennifer Moïsi, Jennifer Verani, Jeremy Keenan, Khalid Zerouali, Maria-Cristina de Cunto Brandileone, Margaret Ip, Md Hasanuzzaman, Nicole Wolter, Noga Givon-Lavi, Özgen Köseoglu Eser, Pak Leung Ho, Patrick E Akpaka, Paul Turner, Paula Gagetti, Peggy-Estelle Tientcheu, Philip E. Carter, Pierra Law, Rachel Benisty, Rebecca Ford, Ron Dagan, Sadia Shakoor, Sanjay Doiphode, Shamala Devi Sekaran, Somporn Srifuengfung, Stephen Obaro, Stuart C Clarke, Tamara Kastrin, Theresa J. Ochoa, Waleria Hryniewicz, Veeraraghavan Balaji and Yulia Urban.

## Transparency Declaration

Dr. Gladstone reports PhD studentship from Pfizer, outside the submitted work; Dr. Lees reports grants from Pfizer, outside the submitted work. Dr. von Gottberg reports grants and other from Pfizer, during the conduct of the study and grants and other from Sanofi, outside the submitted work; Dr. Bentley reports personal fees from Pfizer, personal fees from Merck, outside the submitted work. Professor A. Skoczynska reports grants from Pfizer, assistance to attend scientific meetings and honoraria for lecturing funded from GlaxoSmithKline and Pfizer, and participation in Advisory Board of GlaxoSmithKline and Pfizer, outside the submitted work.

## Disclaimer

The findings and conclusions in this report are those of the authors and do not necessarily represent the official position of the Centers for Disease Control and Prevention.

---

**Figure S1.**
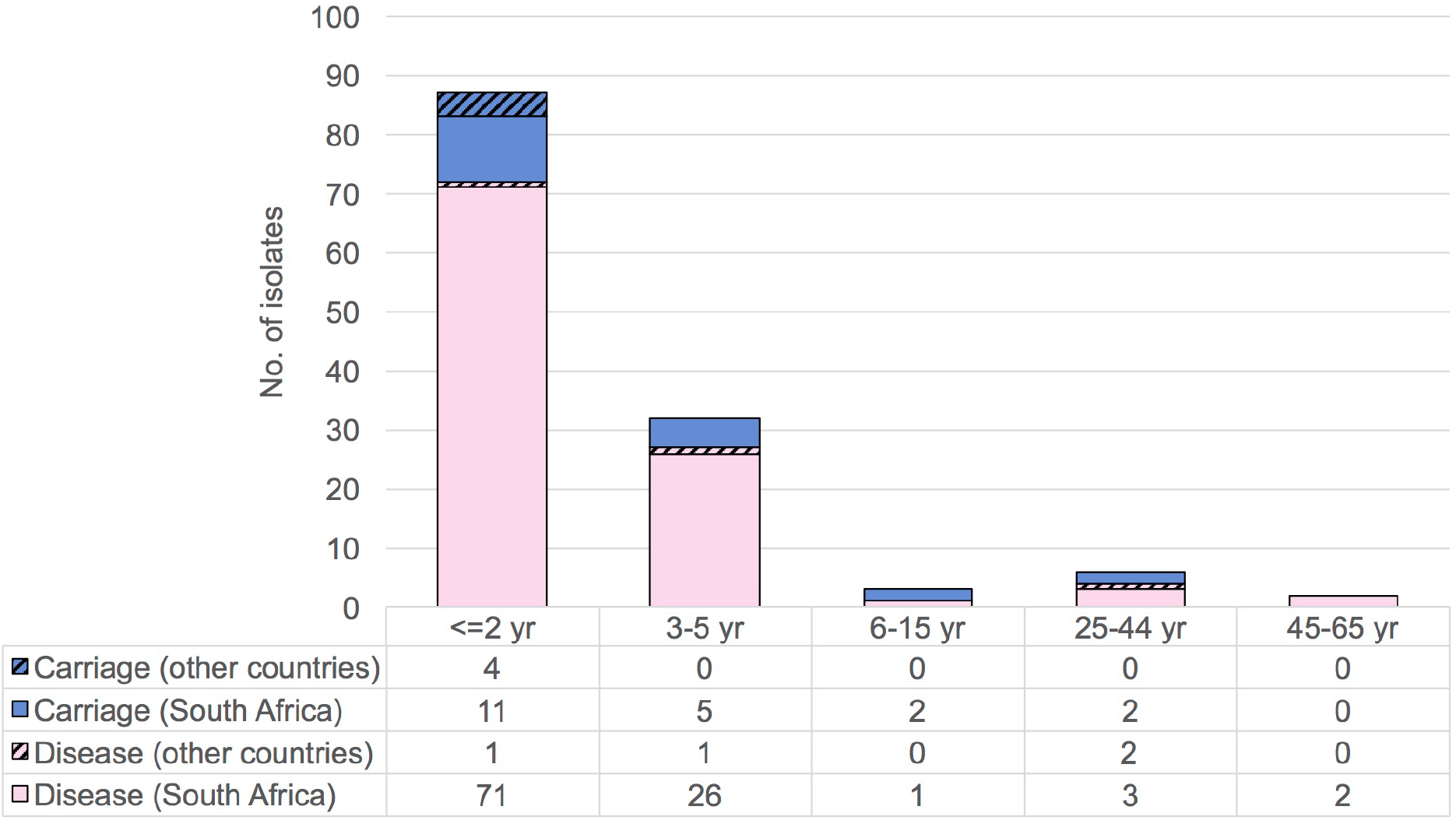
Number of *tet*(S/M)-positive isolates, by age and clinical manifest

**Figure S2.**
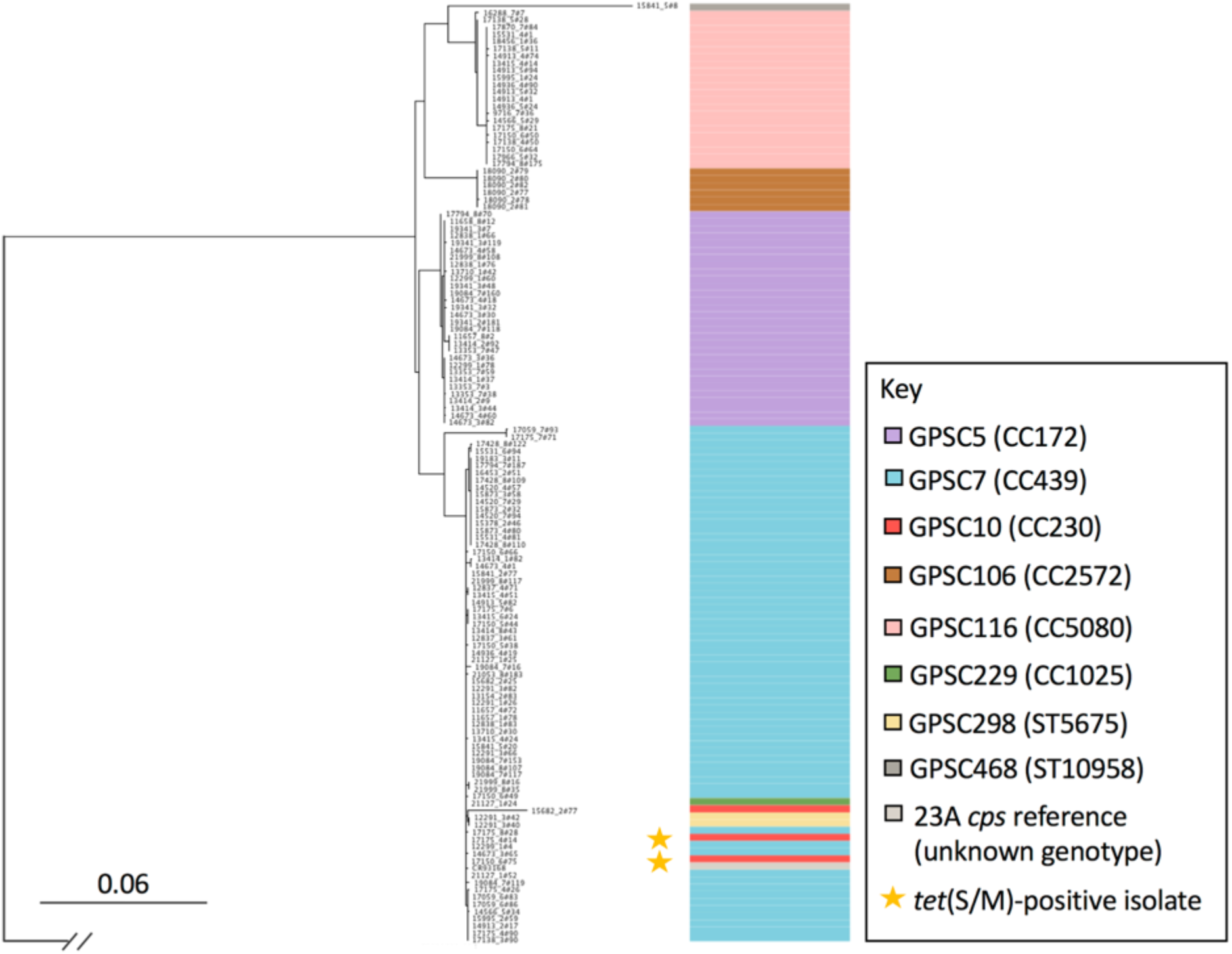
Maximum likelihood phylogenetic tree was constructed using 2,178 SNPs extracted from a 21,296-bp alignment of serotype 23A *cps* locus sequences from the serotype 23A S. pneumoniae isolates (n=130) in the GPS curated dataset. This analysis used the serotype 23F *cps* locus reference sequence (accession number CR931685) as the outgroup on which to root the tree. The serotype 23A *cps* reference sequence (accession number CR931683) was included. The primary clonal complex (CC) or sequence type (ST) associated with Global Pneumococcal Sequence Cluster (GPSC) was indicated in parentheses.

**Figure S3.**
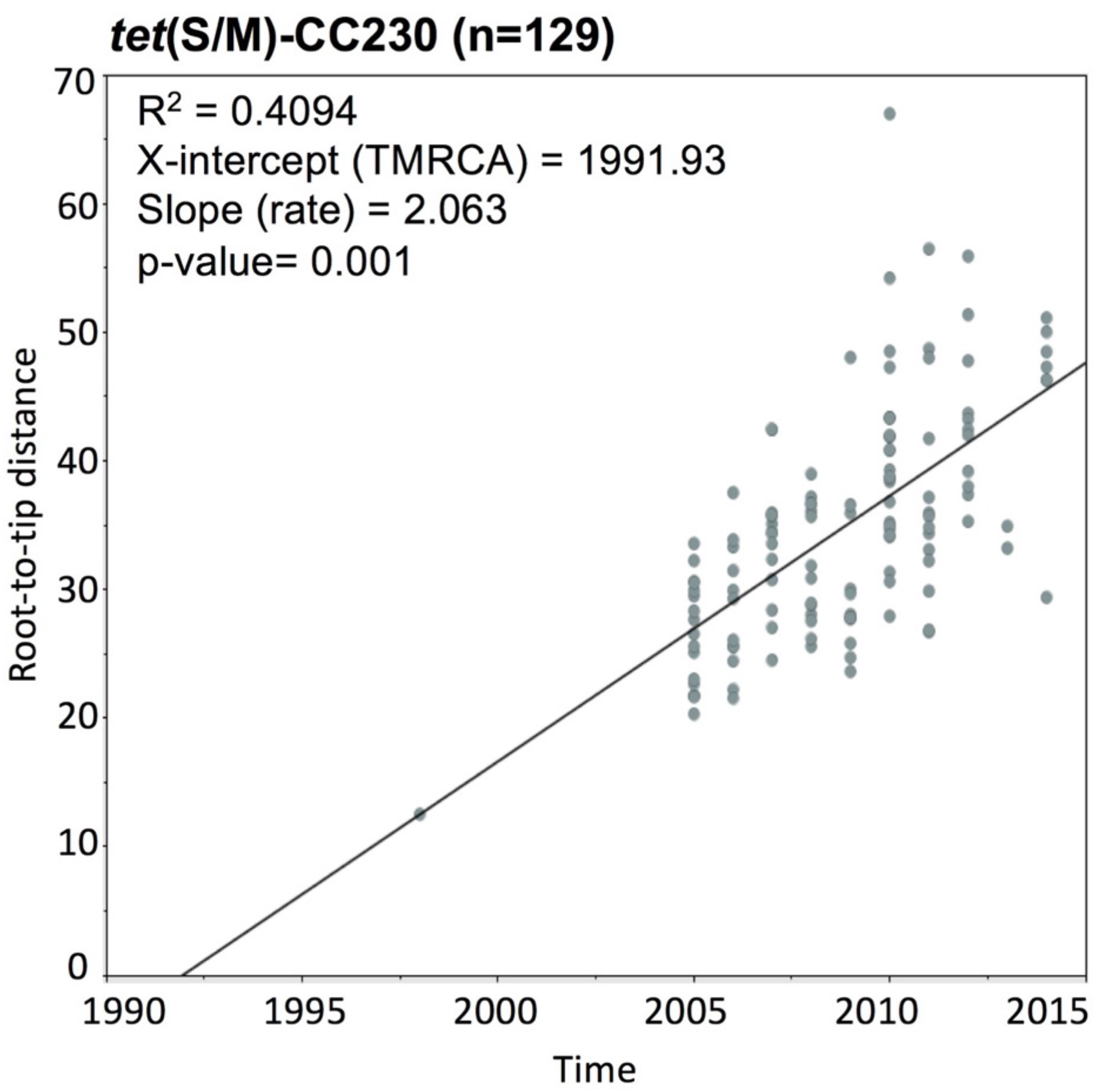
Linear regression of root-to-tip distance against time on *tet*(S/M)-CC230 lineage (n=129) using TempEST v1.5. TempEst detected a significant positive correlation of year of collection with its genetic distance from the root, indicating a signal of a ‘molecular clock’, with which isolates measurably diversifying from their last common ancestor over time.

**Figure S4.**
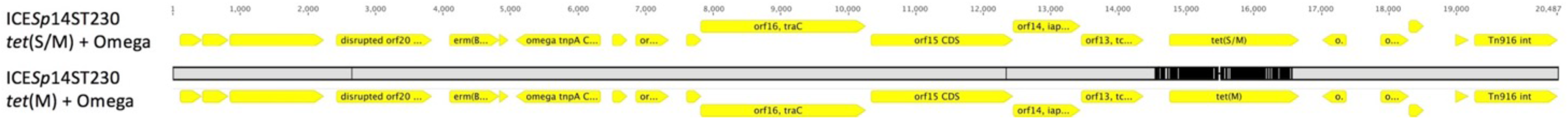
Comparison of Integrative and conjugative element (ICE) carrying *tet*(S/M) and *tet*(M) in clonal complex (CC)230. The yellow arrows indicated protein coding region. The grey band between the sequence indicates BLASTN match and black vertical lines shows the unmatched nucleotide bases.

